# Neuroblastoma formation requires unconventional CD4 T cells and myeloid amino acid metabolism

**DOI:** 10.1101/2021.02.08.430292

**Authors:** Lee-Ann Van de Velde, E. Kaitlynn Allen, Jeremy Chase Crawford, Taylor L. Wilson, Clifford S. Guy, Marion Russier, Leonie Zeitler, Armita Bahrami, David Finkelstein, Stephane Pelletier, Stacey Schultz-Cherry, Paul G. Thomas, Peter J. Murray

## Abstract

By mirroring their function as tissue repair organizers in normal tissues, immune cells regulate tumor growth. To understand the different facets of immune-tumor collaboration through genetics, spatial transcriptomics, and immunological manipulation with non-invasive, longitudinal imaging, we generated a penetrant double oncogene-driven autochthonous model of neuroblastoma. Using spatial transcriptomic analysis, we co-localized CD4^+^ and myeloid populations within the tumor parenchyma, while CD8^+^ T cells and B cells were peripherally dispersed. Depletion of CD4^+^ T cells or CCR2^+^ macrophages, but not B cells, CD8^+^, or NK cells, prevented tumor formation. Tumor CD4^+^ T cells displayed unconventional phenotypes, were clonotypically diverse, and antigen-independent. Within the myeloid fraction, tumor growth required myeloid cells expressing Arginase-1. Overall, our results suggest that arginine-metabolizing myeloid cells conspire with pathogenic CD4^+^ T cells to create permissive conditions for tumor formation, and therefore suggest that these pro-tumorigenic pathways can be disabled by targeting myeloid amino acid metabolism.

## Introduction

Despite the burgeoning interest in tumor immunology and inflammation, and its importance to understanding and treating human cancer, many conceptual and technical limitations are restraining the basic understanding of the immune-tumor interplay and the practical outcomes of new knowledge about cancer. These restrictions encompass widespread use of orthotopic transplantation of tumor cell lines of dubious relevance to human cancer, transplantation to anatomical sites of limited relevance to the natural origins of a given cancer, inability to track and predict immune cell infiltration patterns, and inter-laboratory variance in models. A limitation of mouse cancer models is that they do not accurately reflect the time frame of human malignancies, or the accumulation of mutations. Neuroblastoma (NB) is an ideal ‘laboratory’ to explore the immune-tumor interface. NB arises in the neuroectoderm and mainly forms from the adrenal medulla. The driver oncogenes for NB are well understood (*MYCN, LIN28, ALK*) and form the basis of NB mouse models, which reflect the rate of incidence, location and pathology of human disease (Cazes et al., 2014; Molenaar et al., 2012; Weiss, 1997). The predictable location and incidence of NB allows focus across a window of tumor growth, as opposed to conventional cell line or xenograft systems where a bolus of cells is introduced into an animal.

Immune cells, and especially myeloid cells such as different types of macrophages, dendritic cells, and myeloid-derived suppressor cells, invade tumors from the earliest stages of tumor growth. Large numbers of macrophages and other myeloid cell types, as well as regulatory T cells within tumors, are correlated with poor outcomes in most childhood and adult cancers, spurring a multitude of clinical approaches to target these pathways (Biswas and Mantovani, 2010; Qian and Pollard, 2010). Macrophage infiltration of tumors is inseparable from the process of ‘cancer inflammation’, a ‘hallmark’ of cancer. More specifically, cancer is a type of *non-resolving* inflammation (Murray, 2018; Nathan and Ding, 2010). Other examples of non-resolving inflammation include leprosy, latent tuberculosis, asbestosis, the foreign body reaction to implanted medical devices, and chronic diseases such as Crohn’s Disease and lupus. While the ‘rules’ governing immune cell behavior in *resolving* inflammation in different organs are rapidly emerging, non-resolving inflammation is more complex because the resolving phase of inflammation is absent or aberrant. That the tumor microenvironment is immune ‘suppressive’ is an accepted facet of most cancers because: (i) since tumors are ‘self’ the numbers of antigen-specific lymphocytes are low, and (ii) overcoming tolerance to self-antigens is a key strategy for clinical development of agents to provoke an anti-tumor response. In this regard, attempts to bypass self-tolerance is a key goal of cancer therapy.

Poorly vascularized and hypoxic inflammatory microenvironments are deficient in key nutrients necessary to support expansion of effector immune functions. Examples of these phenomena include infections associated with hypoxic granulomas such as tuberculosis and schistosomiasis (Araújo et al., 2010; Duque-Correa et al., 2014). In these diseases, arginine consumption by myeloid cells is required to suppress tissue destructive T cell proliferation as part of the tissue repair and resolution process (Duque-Correa et al., 2014; Pesce et al., 2009). Within the immune system and especially the myeloid compartment, regulated enzymes metabolize amino acids (Murray, 2016) including Arginase-1 (Arg1), a cytoplasmic hydrolase that metabolizes arginine to ornithine, and Arg2, a closely related arginase to Arg1 that performs the same biochemical reaction, but is restricted to mitochondria. Both enzymes have been reported in multiple tumor types (Bod et al., 2017; Carbonnelle-Puscian et al., 2009) and are under complex regulatory control by cytokines and microbial products. The notion that T cell responses can be suppressed by local myeloid cells expressing amino acid-consuming enzymes is hypothesized to be a component of the regulatory network of ‘infectious tolerance’, where different immune cells propagate to suppress neighboring cells (Andersson et al., 2008; Cobbold et al., 2009).

In cancer, a disease generally associated with aberrant vascularization and hypoxia, both resident myeloid cells and the tumor cells themselves can consume amino acids (Gajewski et al., 2013). Indeed, in pancreatic cancer models, arginine and tryptophan are specifically depleted from the tumor microenvironment relative to all other amino acids (Sullivan et al., 2019). Tumors may have therefore harnessed a natural immunoregulatory program as a mechanism to aid in immune evasion. Arg1 is closely associated with cancer immunosuppression and is a recent clinical target (Arlauckas et al., 2018; Katzenelenbogen et al., 2020; Molgora et al., 2020; Steggerda et al., 2017). The overall goal of targeting myeloid amino acid-consuming enzymes is to enhance local anti-tumor T cell responses, or to overcome a ‘negative’ signal that blocks anti-tumor T cell reactivity. So far, genetic tests of the role of myeloid amino acid-metabolizing enzymes in autochthonous tumors have been limited to a single study where the loss of IDO1 on a KRAS-driven lung cancer model had limited effect in prolonging survival (Smith et al., 2012). We hypothesized that amino acid-dependent pathways are corrupted in the tumor microenvironment, and this phenomenon is linked to the inability of the host to control tumor proliferation and to changes in T cell phenotypic states. To test this, we constructed a genetic system to measure the contribution of a key cancer-associated myeloid amino acid-metabolizing enzyme Arg1 to cancer development and progression.

## Results

### Development of a penetrant neuroblastoma model

NB is the most prevalent solid tumor of children and originates from the sympathetic nervous system that gives rise to the catecholamine-producing adrenal medulla (Matthay et al., 2016). Consequently, NB predominantly arise in the kidney-aorta region. MYCN amplification is the most common genetic abnormality in pediatric NB (Matthay et al., 2016). Tyrosine hydroxylase (TH, the rate-limiting enzyme for catecholamine generation) promoter-driven MYCN transgenic mice are an established model of murine NB which parallel aspects of the childhood disease (Berry et al., 2012a; Molenaar et al., 2012; Weiss, 1997) with tumors exhibiting predictable kinetics, quantifiable by non-invasive means. However, in our vivarium, tumor formation in TH-MYCN mice occurred in less than 10% of screened animals. To improve our understanding of NB biology and intratumoral immunology, we created a highly penetrant autochthonous NB model that builds upon the previous genetic approach. To accelerate tumor formation in the low penetrance TH-MYCN background, we introduced point mutations in *Alk*, a tyrosine kinase oncogene, which, after MYCN, is the most commonly associated gene with human NB. Two gain-of-function mutations were created: *Alk*^F1178L^ (equivalent to human *ALK*^F1174L^, which is the predominant *ALK* mutation in human NB (Berry et al., 2012b)), and *Alk*^Y1282S^ (equivalent to human *ALK*^Y1278S^, also found in human NB) (Janoueix-Lerosey et al., 2008; Pugh et al., 2013). (**Fig. 1a, Extended Data Fig. 1a, b**). The F1178L or Y1282S mutations were crossed into the TH-MYCN background in heterozygosity. We longitudinally tracked NB formation and growth by ultrasound imaging (**Extended Data Table 1**) within a design appropriate to endpoint comparison of individual animals across time (**Fig. 1b**). Mice with animal welfare issues unrelated to a tumor were excluded **(Extended Data Table 2**). Mice with *Alk*^F1178L^ on a TH-MYCN background developed NB with an incidence of ∼50% within the study window, and approximately 40% of mice achieved a study endpoint within the study period. The *Alk*^Y1282S^ mutation on the TH-MYCN background produced approximately half the tumor incidence and survival rate of the *Alk*^F1174L^ mutation (**Fig. 1c**). Consistent with other studies (Berry et al., 2012a; Durbin et al., 2018; Heukamp et al., 2012), we found that the transcriptomes of the tumor samples were concordant with human NB and distinct from other childhood solid tumors (**Fig.1d, Extended Data Table 3 and 4**). We observed significant expression of the current drug targets GD2 **(Fig. 1e)** and B7-H3 (CD276) **(Fig. 1f)**, and low amounts of PD-L1 and PD-L2 (**Fig. 1g**), replicating expression patterns seen in human NB (Aoki et al., 2016; Castriconi et al., 2004; Majzner et al., 2017; Schulz et al., 1984) and demonstrating the utility of this animal model as a pre-clinical tool. For all remaining longitudinal studies, we used the high penetrance *Alk*^F1178L^; TH-MYCN mice to investigate how manipulating amino acid metabolic pathways contributes to tumor growth. Unlike the widely used xenograft transplantation models, our autochthonous, genetics-based model system allows for precise identification of the conditions in which malignancy is established and permits detailed analysis of the earliest stages of the anti-tumor immune response.

**Fig. 1.**
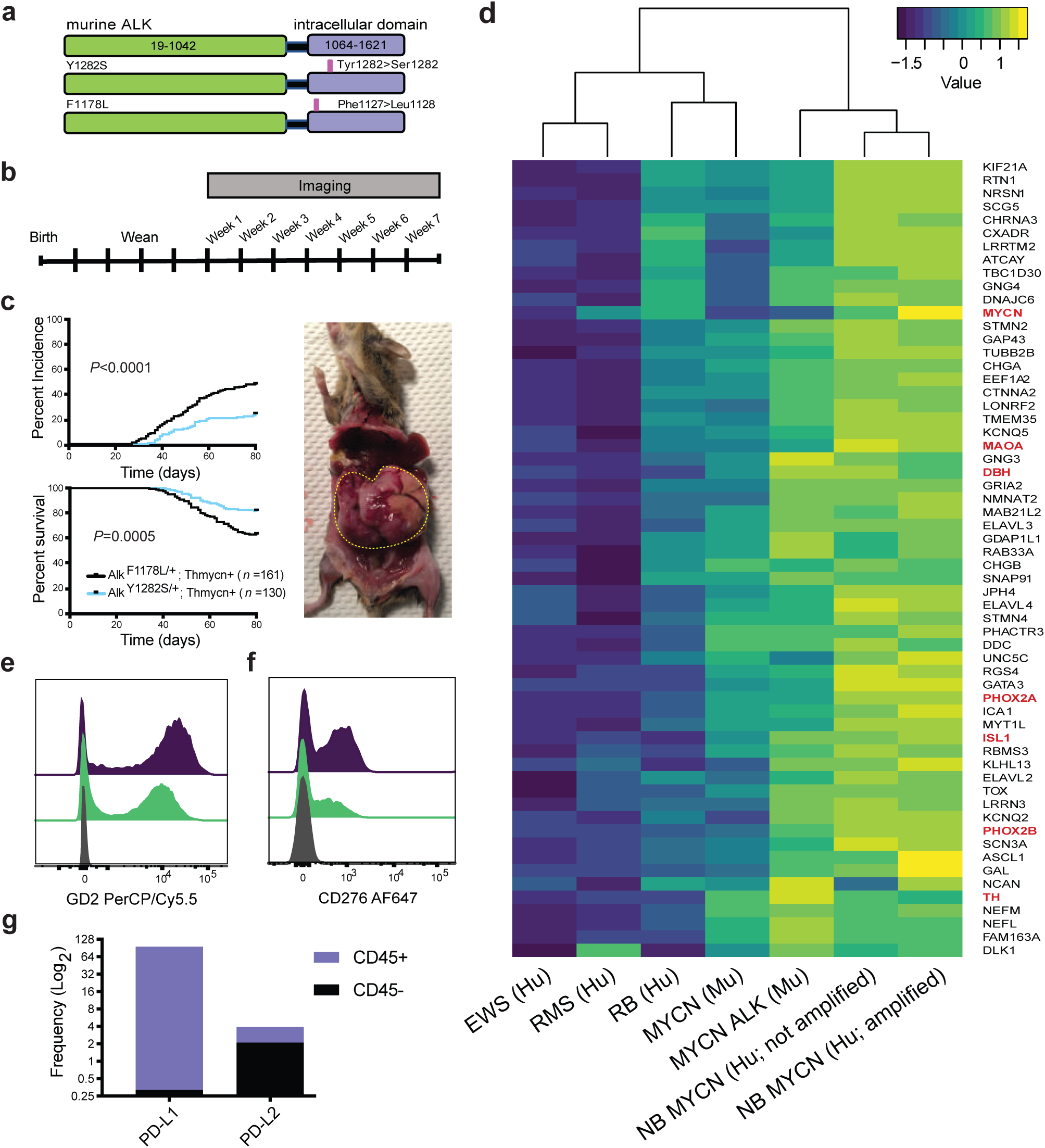
Point mutations in the endogenous *Alk* locus and NB formation. **a**, Diagram of the two ALK mutations introduced into the endogenous *Alk* locus. **b**, Experimental design of the longitudinal study for animals reported herein. **c**, Incidence and survival curves comparing tumor formation of *Alk*^F1178L^; TH-MYCN or *Alk*^Y1282S^; TH-MYCN mice. Representative image of an *Alk*^F1178L^; TH-MYCN mouse with a large abdominal NB. **d**, Clustered heat map of ‘signature’ gene expression comparing murine NB models and other human solid tumors. **e, f**, GD2 and CD276 (B7-H3) expression in tumor cells. **g**, Frequency of PD-L1 and PD-L2 expressing cells in CD45^+^ or CD45^-^ intra-tumoral populations. In **c**, data were analyzed by the Log-rank test. Criteria for scoring tumor incidence and inclusion and exclusion criteria are described in Methods. In **d**, the complete transcriptional signature is available in **Extended Data Table 3** and available in full using Accession #GSE12460, GSE13136, GSE16237, GSE37372, GSE29683, E-TABM-1202, GSE27516, GSE98763, GSE27516.

### Single-cell sequencing identified transcripts encoding amino acid metabolizing enzymes

We next sought to characterize the immune infiltrate of our NB tumor model using single-cell transcriptomics. We collected single-cell gene expression (scGEX) data on our *Alk*^F1178L^; TH-MYCN tumors by isolating tumor cells and parenchymal immune cells, which were identified by injecting an anti-CD45 antibody intravenously prior to euthanizing the animals. After tumor processing, we stained with a CD45 antibody conjugated with a different fluorophore, allowing us to segregate vascular and parenchymal immune cells. Parenchymal immune cells and tumor were mixed at equal ratios for scGEX profiling (**Fig. 2a, Extended Data Figure 1c)**.

**Fig. 2.**
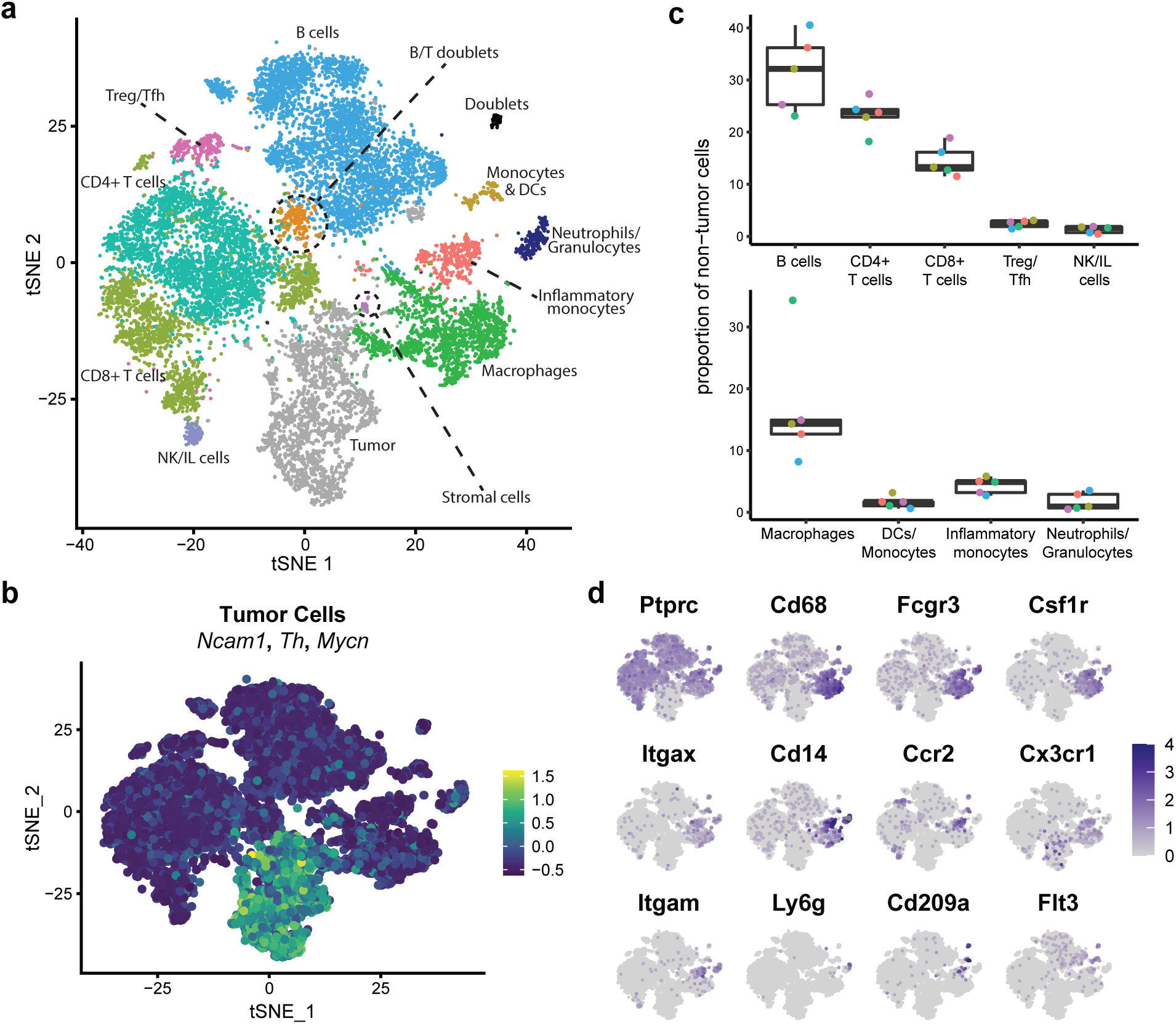
Single-cell gene expression reveals tumor infiltrating immune populations. **a**, t-SNE dimensionality reduction of mouse *Alk*^F1178L^; TH-MYCN CD45^-^ and CD45^+^ tumor cells based on single-cell gene expression (scGEX) data, with cell clusters identified based on marker gene expression (**Extended Data Fig. 1c**). **b**, Feature plot of tumor cells indicated by module scoring of of *Ncam1, Th*, and *Mycn* expression. **c**, Box plots representing the proportion of cell types among the tumor-infiltrating CD45^+^ cells using the clusters identified in the scGEX analysis. Colors indicate replicate tumor identity within and across plots. **d**, Feature plots showing expression of gene markers of specific myeloid populations found in the tumor sample.

Tumor cells were identified as those expressing known NB gene hallmarks including *Mycn, Th*, and *Ncam1*, **(Fig. 2b, Extended Data Fig. 1d)** with no expression of *Ptprc*. This signature identified three tumor clusters which comprised 16.9% of the total cells captured. Unsurprisingly, the main genes enriched in these clusters compared to non-tumor immune cells included many known to be associated with tumor biology, including *Tuba1a* (Wang et al., 2020), *Stmn2* (Byrne et al., 2014), *Uchl1* (Kwan et al., 2020), and *Tubb3* (Person et al., 2017)

**(Extended Data Fig. 1d)**. Known neuronal genes were upregulated in the tumor clusters, including *Pirt* (Tang et al., 2013), *Tmeff2* (Masood et al., 2020), and *Dst* (Dalpé et al., 1998), which have been previously identified as upregulated genes in NB tumors (Brady et al., 2020) **(Extended Data Fig. 1d**). For instance, *Pirt* is a gene specifically expressed in peripheral sensory neurons that is differentially methylated in varying stages of NB tumor progression (Tang et al., 2013). Overall, the transcriptional profiling of tumor cells substantiated the complex nature of NB, with the observation of neuronal signatures and various tumor biology signatures including growth, adhesion, proliferation, invasion, and metastasis.

Among the non-tumor immune cells, B cells dominated the cellular enumeration (32%), followed by CD4^+^ T (26%) cells, and CD8^+^ T (14.5%) cells (**Fig. 2c, Extended Data Fig. 1e)**. Within the innate compartment, discrete populations of macrophages and monocytes encompassing various states of activation and development were present, with smaller populations of neutrophils, granulocytes, NK cells, and innate-like lymphocytes also observed. Among the myeloid populations, we distinguished macrophages, inflammatory monocytes, neutrophils, and dendritic cells/immature monocytes based on gene expression (**Fig. 2d**). The largest component of this compartment were macrophages, which made up over 13% of the total immune cell population, with only a few percent of cells in the other compartments.

While scGEX analyses are useful for resolving the cellular heterogeneity of the tumor, the spatial organization of these cells can significantly impact their role within the tumor microenvironment. To explore this, we performed spatial transcriptomic analysis on sections from multiple tumors. We leveraged the scGEX data by integrating the single-cell and spatial molecular signals and identifying overlaps in expression patterns. Unsurprisingly, the presence of the scGEX tumor signature was nearly ubiquitous throughout the spatial transcriptomics data, though notably absent in large areas of presumptive vasculature. The distinct transcriptional clusters identified within this broader tumor signature appeared to correspond with spatial features within the tumor environment, where the most prevalent tumor scGEX cluster (cluster 3, **Extended Data Fig. 1c**) was most enriched at the tumor core, whereas the second most prevalent tumor cluster (cluster 7) was more spatially dispersed (**Fig. 3a**), indicating distinct tumor microenvironments that correspond with density and immune infiltration. Further analysis to compare tumor clusters 3 and 7 demonstrated that cluster 3 was more enriched in neural-like genes **(Fig. 3b)**. Though both tumor clusters were enriched with many cancer-associated genes, cluster 3 in general expressed numerous genes known to be involved in multiple facets of tumor biology at higher levels than cluster 7 **(Fig. 3b; Extended Data Fig. 1d**).

**Fig 3.**
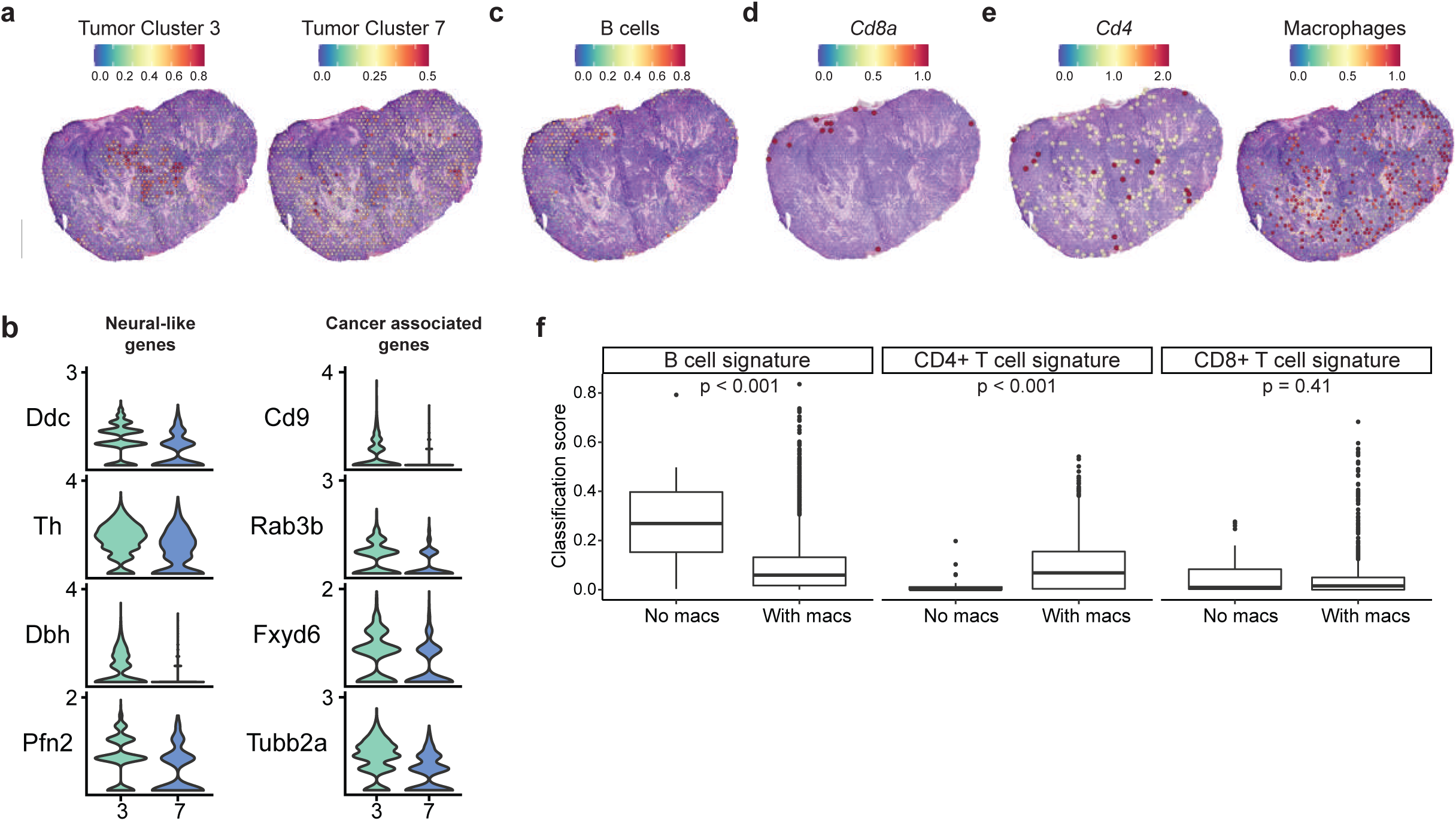
NB landscape is characterized by specific spatial patterns of immune and tumor cell subsets. **a**, Spatial transcriptomics feature plots from a representative section (A1) where expression of individual genes is overlaid on top of tumor section. Capture area points are enlarged for ease of visualization and are transparent for values below 0.1 on the accompanying color scale. Tumor cluster signatures represent classification scores derived from integration of single-cell gene expression data with spatial transcriptomics data. **b**, Violin plots of genes enriched in tumor cluster 3 compared to tumor cluster 7, including neural-like genes and cancer-associated genes. **c, d, e**, Spatial transcriptomics feature plots from a representative NB section (A1) for B cells, CD8^+^ T cells, CD4^+^ T cells, and macrophages. *Cd8a* and *Cd4* values are from section-specific log normalization. **f**, Box plots showing scGEX signature classification scores from spatial transcriptomics section A1 for capture spots with a macrophage classification score of 0 (*no macs*) versus spots with a positive macrophage classification score (*with macs*). Statistical comparisons were made with Wilcoxon Rank Sum tests.

We next examined the distribution of the innate and adaptive immune cell subsets in the tumor. Although B cells constituted a large proportion of the lymphocytes identified in the scGEX analyses, their corresponding expression signature was notably absent from the inner areas of the tumor and instead were enriched around the tumor periphery (**Fig. 3c**). Similarly, CD8 T cells were not evenly distributed within the tumor, instead mostly accruing in peripheral areas (**Fig.3d**). In contrast, both CD4^+^ T cells and macrophages were widely distributed throughout the tumor (**Fig. 3e**). Notably, CD4^+^ T cell classification scores were significantly higher among areas with a positive macrophage classification score **(Fig. 3f)**, indicating colocalization between these two cell subsets.

### CD4^+^ T cells are pathogenic in neuroblastoma

Given the significant presence of adaptive immune cells in the tumor parenchyma, we next wanted to determine their contribution to tumor restraint or growth. To address the role of both B cells and T cells, we first crossed *Alk*^F1178L^; TH-MYCN mice with *Rag1*^-/-^ animals to generate a strain lacking B and T cells from birth. Strikingly, we have never observed a single tumor in this line, indicating that that an intact adaptive immune compartment is crucial for NB formation in this model (**Fig 4a**). We therefore undertook a series of experiments to identify the effectors of this pro-tumorigenic response. The role of B lymphocytes in the anti-tumor immune response to solid tumors remains poorly understood, with both anti- and pro-tumor roles reported (Helmink et al., 2020; Petitprez et al., 2020; Sharonov et al., 2020), and no studies to date have systematically investigated the role of B cells in NB. To determine the overall contribution of B cells to tumorigenesis in our model, we treated *Alk*^F1178L^; TH-MYCN mice with dual anti-CD19 and anti-B220 depleting antibodies. We observed no significant differences in tumor incidence or survival rates (**Fig. 4b**), suggesting that B lymphocytes are not required for tumor formation.

**Fig. 4.**
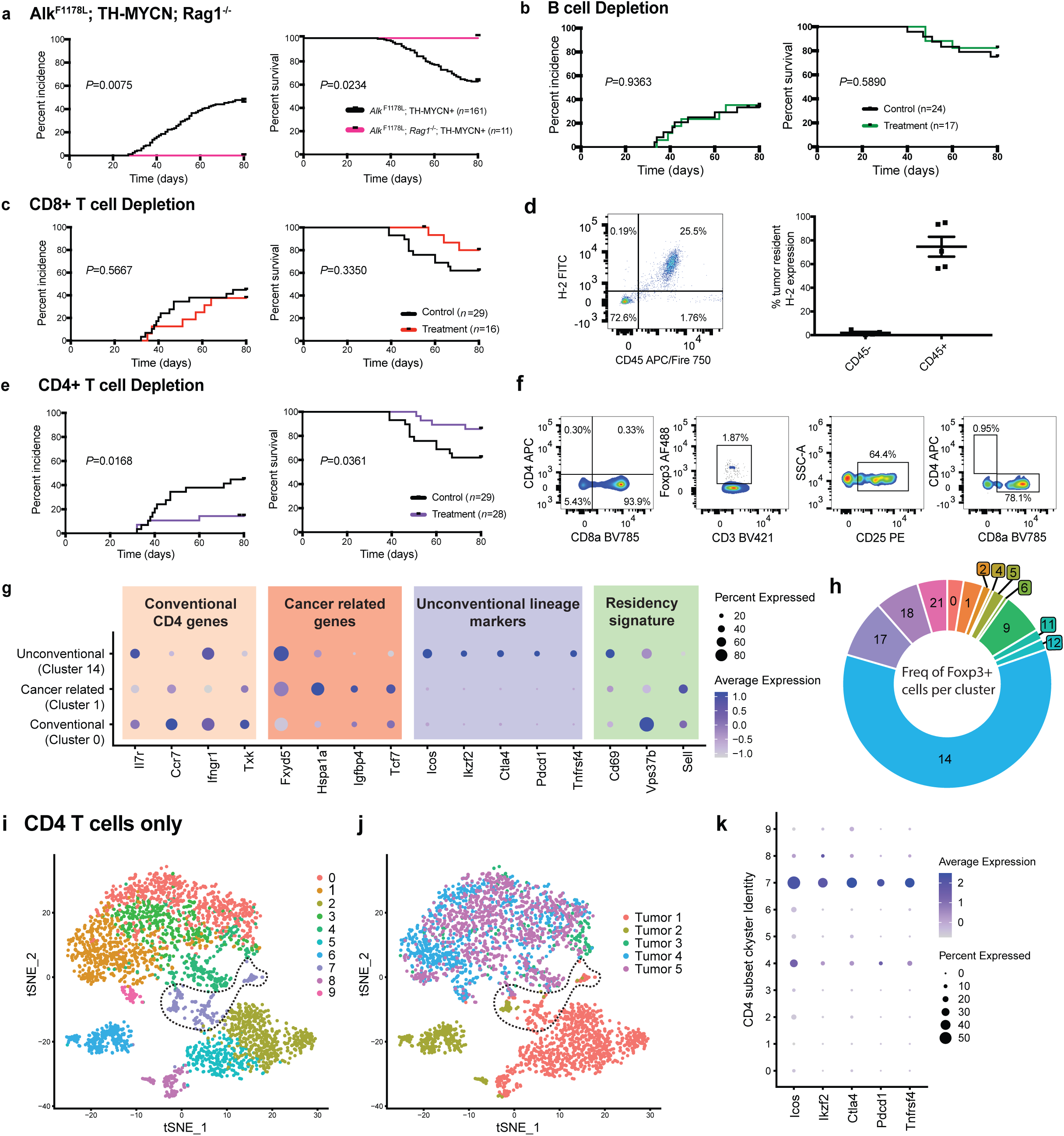
CD4^+^, but not CD8^+^ or B cells, are pathogenic in *Alk*^F1178L^; TH-MYCN NB. **a**, Incidence and survival curves comparing tumor formation between *Alk*^F1178L^; *Rag1*^-/-^, TH-MYCN mice and *Alk*^F1178L^; TH-MYCN mice. **b**, Incidence and survival curves of *Alk*^F1178L^; TH-MYCN mice treated with dual anti-CD19 and anti-B220 antibodies or control. **c**, Incidence and survival curves in *Alk*^F1178L^; TH-MYCN treated with anti-CD8 depleting antibody or control. **d**, Flow cytometric analysis of MHC class I expression on intra-tumoral CD45^+^ and CD45^-^ populations. **e**, Incidence and survival curves in *Alk*^F1178L^; TH-MYCN mice treated with anti-CD4 depleting antibody or control. **f**, Successful depletion of CD4^+^ T cells in the tumor (1^st^ panel). Presence of Foxp3^+^ CD8^+^ cells in *Alk*^F1178L^; TH-MYCN mice treated with anti-CD4 depleting antibody. Data are representative of *n*=2 mice. **g**, Dot plot of genes enriched across the CD4^+^ T cell clusters in these tumor samples with genes binned in functional groups “Conventional CD4 genes,” “Cancer-related,” “Unconventional lineage markers,” and “Residency genes.” Color corresponds to expression of each gene relative to the average among the four focal populations, and the size of the dot represents the proportion of cells from the cluster expressing each gene. **h**, Donut plot of the frequency of Foxp3^+^ cells within each cluster. Clusters with 0% cells expressing Foxp3 are not shown. **i**, t-SNE of only CD4^+^ T cells subsetted from the full dataset, with sub-clusters identified based on marker gene expression. Dotted line indicates CD4 cluster 7. **j**, CD4 t-SNE from **i**, with colors indicating the originating tumor. Dotted line indicates CD4 cluster 7. **k**, Dot plot of “Unconventional lineage markers.” In **a, b, c**, and **e**, data were analyzed by the Log-rank test.

We next measured the effect of depleting CD8^+^ T cells, though in our spatial transcriptomic data these cells did not appear to be well-infiltrated in the tumor. CD8^+^ T cells were dispensable for tumor formation, with depleted animals demonstrating similar rates of tumor development as control animals (**Fig. 4c)**. CD8^+^ T cells exert their anti-tumor function in part by recognizing tumor-associated antigens on MHC class I. scGEX and spatial transcriptomic data indicated that the tumor cells failed to express significant levels of β2-microglobulin (**Extended Data Fig. 2a**). Similarly, human pediatric NB express extremely low levels of β2-microglobulin, especially when compared to other tumor types (**Extended Data Fig. 2b**). Staining for MHC class I expression, we found that the tumor cells were largely class I negative, confirming our transcriptional data (**Fig. 4d**) and suggesting that NB downregulate these recognition molecules to blunt an effective tumor-specific cytotoxic CD8^+^ T cell response. Although MHC class I-independent tumor control can occur through NK cell mediated cytotoxic activity, the depletion of NK cells also had no effect on tumor incidence or survival (**Extended Data Fig. 2c**).

As both B cells and CD8^+^ T cells were dispensable for tumor formation, we turned to CD4^+^ T cells as the likely candidate for the established pro-tumorigenic effect of adaptive immunity. We depleted total CD4^+^ cells via monoclonal antibody treatment in *Alk*^F1178L^; TH-MYCN mice starting one week prior to study enrollment. Strikingly, CD4^+^ depletion, which depletes both conventional and regulatory CD4^+^ populations, prevented tumor formation **(Fig. 4e)**. We noted that a very small number of CD4^+^-depleted animals still developed tumors. In two of these mice, we analyzed their tumor microenvironment to confirm successful CD4 depletion and observe the phenotype of any residual CD4^+^ T cells that might account for the tumor formation. While no CD4^+^ cells were observed in the blood or spleen, an exceptionally small number were observed within the tumor and appeared to be conventional CD4^+^ T cells, with no expression of Foxp3 (which would be indicative of a regulatory phenotype) (Fontenot et al., 2003) (**Fig. 4f)**. Surprisingly, a population of Foxp3^+^ CD8^+^ cells were enriched in these tumors. These populations were not observed in non-depleted mice **(Extended Fig. 3**), suggesting that this CD8^+^ regulatory phenotype emerges as a compensatory mechanism in the absence of CD4^+^ T cells, and that, at minimum, T cells with the capacity for regulatory activity are required for tumor formation.

The spatial transcriptomic data showing infiltration of CD4^+^ T cells along with these depletion data establish a critical role for CD4^+^ T cells in promoting tumor growth. This prompted us to define the features of these cells within the tumor. In our scGEX analysis, three clusters of CD4^+^ T cells were identified. Analysis of distinguishing genes allowed us to characterize them as: “conventional”, expressing standard markers of naïve and memory T cells and tissue residency (including *Il7r, Ccr7, Ifngr1, Vps37b*); “cancer-related”, expressing genes known to be enriched in cancers, including *Fxyd5, Hspa1a, Tcf7*, and *Igfp4*; and “unconventional”, expressing an array of genes representative of unconventional T cell populations (including innate-like T cells and Tregs), such as *Icos, Ikzf2, Ctla4, Pcdc1*, and *Tnfrsf4* (**Fig. 4g**). Importantly, the “unconventional” cluster was the only cluster with appreciable *Foxp3* expression (**Fig. 4h**).

Our scGEX analysis was based on a pooled dataset from multiple tumors. Given the potential for intertumoral heterogeneity, we analyzed the scGEX clustering of the CD4^+^ T cells only, identifying nine subclusters by k-means clustering (**Fig. 4i**). Using this finer discrimination, we observed one cluster that was common to all tumors (**Fig. 4j**), which displayed an “unconventional” gene expression pattern (**Fig. 4k**). Given the absolute requirement of CD4^+^ cells for tumor formation, these data suggest that CD4^+^ cells with a distinct “unconventional” phenotype may be responsible for this tumor-promoting effect.

CD4^+^ T cells, like all T cells, are defined by the presence of a T cell receptor conferring specificity to an epitope (usually peptide in complex with MHC class II). However, T cells can also perform more “innate” like functions, analogous to TCR-deficient innate lymphocytes, in a TCR-independent manner. Along with nuclear expression of the transcription factor Foxp3, CD4^+^ T cells with a regulatory phenotype highly express IL2rα, or CD25, on their cell surface (Sakaguchi et al., 1995, 2001; Shevach et al., 2001). Since T cells with regulatory potential appear to be essential to tumorigenesis, we examined whether the CD4^+^ CD25+ T cells in our mouse NB shared an epitope-specificity. We performed single-cell TCR sequencing on parenchymal CD4^+^ CD25+ T cells isolated from multiple animals. We and others (Dash et al., 2017) have previously shown that T cells that share a specificity for an epitope target usually carry TCRs with convergent motifs across individuals. Applying the TCRdist algorithm to detect the presence of such motifs, we found that there were no consistent motifs either within or between tumors that would indicate clonal expansion or shared specificity **(Fig. 5a, b**).

**Fig. 5.**
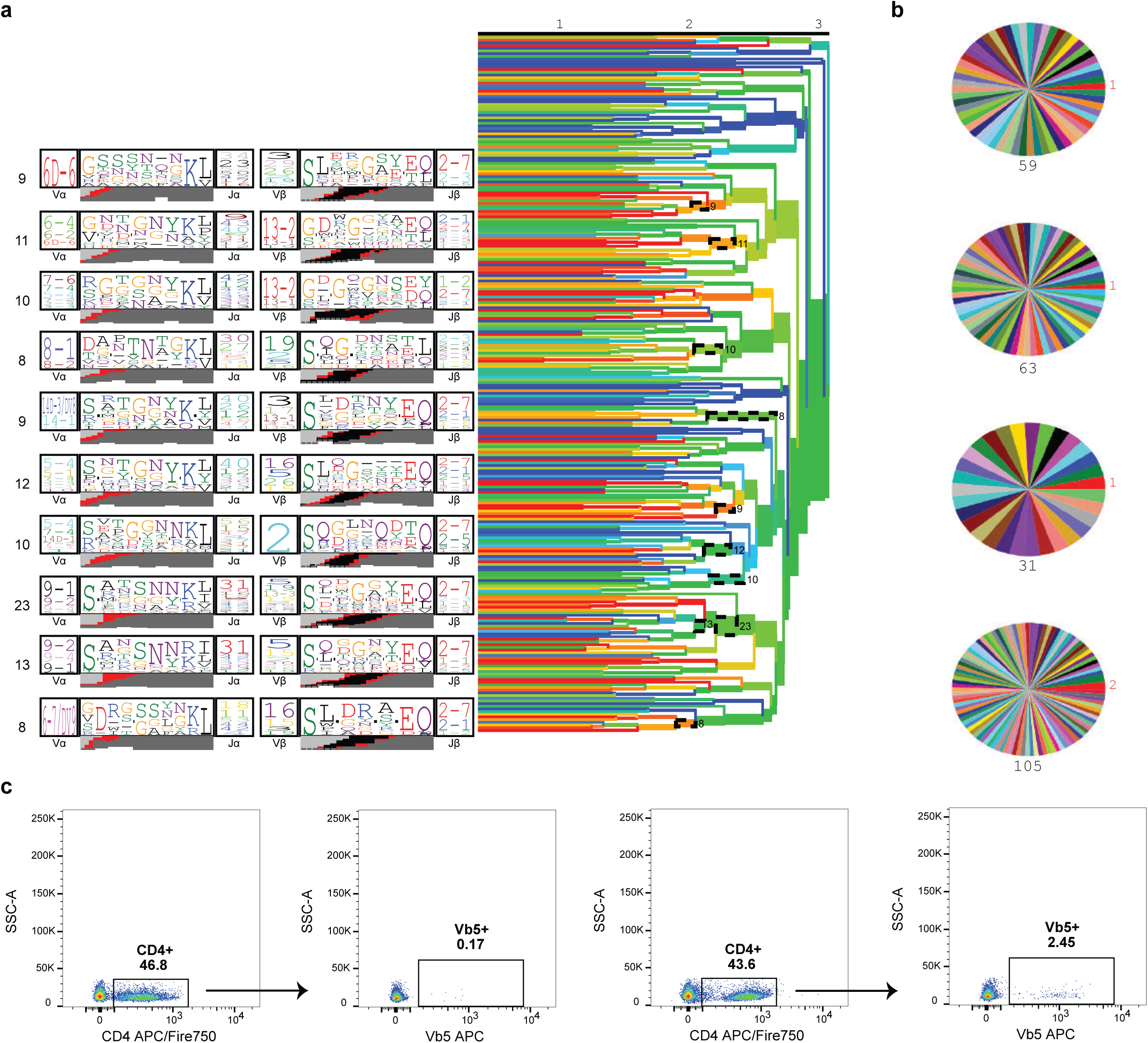
Tumor-resident CD4^+^ CD25^+^ cells are not antigen specific. **a**, CD4^+^ CD25^+^ cells were isolated from *Alk*^F1178L^; TH-MYCN mice for TCR repertoire analysis. TCRdist trees (right) with dashed ellipses indicating groups of similar TCRs, or neighbors, as a cluster. Representative TCR logos (left) depict most highly used V- and J-genes, CDR3 amino acid sequences, and predicted VDJ rearrangement of each cluster. (*n*=4 mice, 262 TCR clones). **b**, Clone pie charts for four independent mice. Each wedge represents a unique TCR clone. The size of top clone is in red, and total number of sequences are in black. **c**, CD4 cells migrate to the tumor in an antigen-independent manner. *Alk*^F1178L^; TH-MYCN mice were intraveneously injected with 5 x 10^6^ CD4^+^ cells from OT-II mice prior to tumor formation. Tumors from mice receiving OT-II donor cells show increased numbers of Vβ5^+^ CD4 cells.

To further confirm that the tumor infiltrating CD4^+^ T cells were acting in an antigen-independent manner, we transferred ovalbumin-specific OT-II CD4^+^ T cells to mice one week prior to study enrollment and the onset of detectable tumors. We then analyzed whether these cells would be recruited into the tumor parenchyma, despite the animals lacking expression of cognate antigen. Indeed, we observed an increased frequency of intratumoral CD4^+^ cells expressing Vβ5 TCR from mice transferred with OT-II CD4^+^ T cells compared to wild-type mice (**Fig. 5c**), suggesting that the phenotype of tumor-resident CD4^+^ cells is obtained through a mechanism that does not require a specific antigen.

### Bone marrow-derived CCR2^+^ myeloid cells drive neuroblastoma

Our spatial transcriptomic data indicated that in CD4 T cells, macrophage and monocyte clusters were the only immune cell types well-infiltrated throughout the tumor. We considered the co-occurrence of the distinct immune populations across all spatial transcriptomics sections. Based on probabilistic classification scoring of 6,729 spatial transcriptomics gene expression spots, we found a clear enrichment in the co-occurrence of moderate CD4^+^ T cell and macrophage classification scores that was not present for other cell subsets (**Extended Data Fig. 4a**). In solid tumors, most macrophages originate from monocytes, whose egress from the bone marrow is largely controlled by the chemokine CCL2 and its receptor CCR2 (Qian et al., 2011). We found *Alk*^F1178L^; TH-MYCN; *Ccr2*^-/-^ mice had >80% depletion of circulating monocytes compared to cognate controls (**Extended Data Fig. 4b**), and these mice had a trend to reduced tumor incidence and significantly increased survival compared to *Alk*^F1178L^; TH-MYCN mice (**Fig. 6a**). In the tumors arising in *Alk*^F1178L^; TH-MYCN; *Ccr2*^-/-^ mice, we observed a relative depletion of monocytes and macrophages compared to controls, as expected (**Extended Data Fig. 4c, d**). Consistent with the “leaky” phenotype of *Ccr2*^-/-^ mice, it is possible some residual monocytes can seed tumors and potentially interact with embryonic macrophages present in the adrenal gland, but nevertheless, these data demonstrate that bone marrow-derived CCR2-dependent monocytes and macrophages contribute to NB development and growth.

**Fig. 6.**
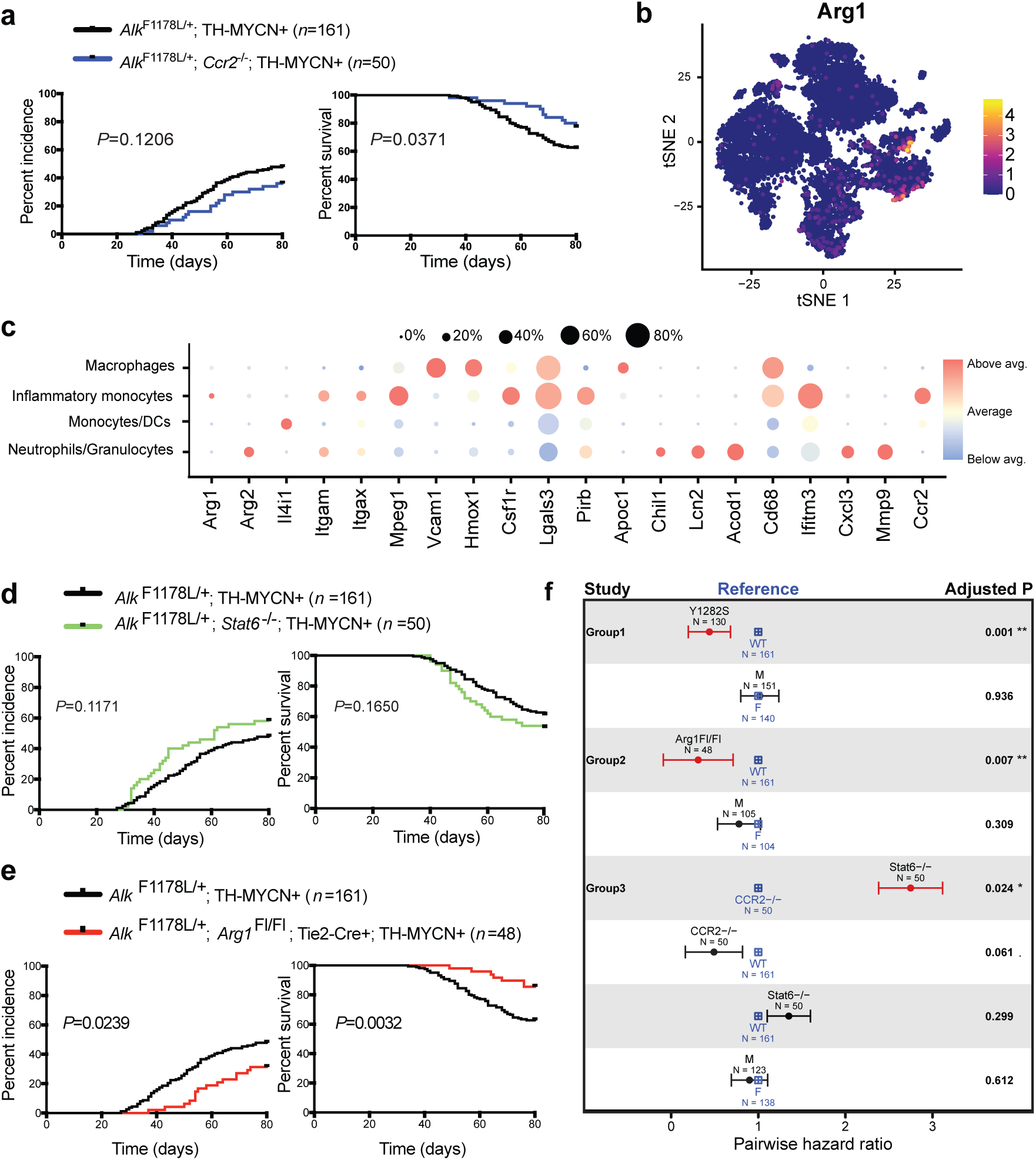
Myeloid Arg1 is pathogenic in NB. **a**, Incidence and survival curves comparing tumor formation of *Alk*^F1178L^; TH-MYCN; *Ccr2*^-/-^ mice. **b**, Feature plots showing cell-specific expression of *Arg1*.**c**, Dot plot of enzymes and known markers of monocyte, macrophage, and granulocyte populations. Color corresponds to expression of each gene relative to the average among the four focal populations, and the size of the dot represents the proportion of cells from the cluster expressing each gene. **d**, Incidence and survival curves comparing tumor formation between *Alk*^F1178L^; TH-MYCN; *Stat6*^-/-^mice and *Alk*^F1178L^; TH-MYCN mice. **e**, Incidence and survival curves comparing tumor formation of *Alk*^F1178L^; TH-MYCN mice lacking *Arg1* in all hematopoietic cells. **f**, Pairwise hazard ratios as determined by CoxPH regression modeling. Reported *P* values are adjusted for multiple comparisons. M=male, F=female. Error bars represent the standard error of the coefficients. In **a, d**, and **e**, data were analyzed by the Log-rank test.

### Myeloid Arg1 is a tumor driver

CCR2-dependent monocytes and macrophages that reside within tumor microenvironments have diverse functions that provide growth and survival advantages to the tumor cells, and importantly, suppress anti-tumor immune responses. A major means by which this occurs is via the production of the amino acid-metabolizing enzyme Arginase-1. Arg1 is reported to be expressed in a variety of tumor environments (Arlauckas et al., 2018; Katzenelenbogen et al., 2020; Molgora et al., 2020), both by resident myeloid cells and the tumor cells themselves (Jang et al., 2018; Krzystek-Korpacka et al., 2020; Singh et al.). Arg1 depletes L-arginine from the local environment and hydrolyzes it to ornithine and urea. T cells are exquisitely sensitive to the deprivation of L-arginine, resulting in proliferative arrest (Van de Velde et al., 2017) and, in the case of CD4^+^ cells, the promotion of Th2 and Treg phenotypes (Castellano and Molinier-Frenkel, 2020). Our scGEX data revealed that in our model of NB, *Arg1* expression was restricted to monocytic cells only, with minimal levels detected within the tumor cells (**Fig. 6b**,**c**). Similarly, immunohistochemical analysis of human NB revealed Arg1 expression only in interspersed myeloid-like cells that overlapped partly with CD68 and CD163 staining **(Extended Data Fig. 5a)**. Our data thus far demonstrate a spatial relationship between resident CCR2-dependent myeloid cells and non-antigen specific CD4^+^ cells, and that both populations are required for tumor formation. Considering the conserved “unconventional” phenotype in intratumoral CD4^+^ cells, characterized by the expression of a suite of genes associated with suppressive activity, and prior links between amino acid metabolism and Treg induction (Cobbold et al., 2009; Murray, 2016; Qin et al., 1993), we sought to investigate the role of Arg1 within our model.

Within the tumor microenvironment, Arg1 expression can be linked to multiple functional groups of myeloid cells. Tissue-infiltrating monocytes can differentiate into macrophages with distinct effector phenotype, including broadly defined ‘M1’ and ‘M2’-like cells. Most tumor-associated monocytes and macrophages (TAM) are considered to be M2-like, which are key components of tissue repair and wound healing, and actively promote tumor growth through multiple mechanisms (Biswas and Mantovani, 2010; Eming et al., 2017). The canonical pathway for M2 macrophage polarization employs STAT6 downstream of the IL-4 and IL-13 receptors, and in coordination with KLF-4, induces the expression of M2-related genes, including *Arg1* (Murray, 2017), which potently suppress the anti-tumor immune response. To test whether the monocytic contribution to tumor formation depended on these M2 pathways, we crossed our *Alk*^F1178L^; TH-MYCN mice to mice lacking *Stat6*. We found no evidence that IL-4/IL-13 M2 macrophages were drivers of disease, as tumor formation in *Alk*^F1178L^; TH-MYCN; *Stat6*^-/-^ were similar to their cognate controls (**Fig. 6d)**. Thus, while monocyte-differentiated macrophages contributed to NB formation, their effects were not mediated through Stat6-dependent, M2-dependent pathways. These data are consistent with previous studies demonstrating that Arg1 can be induced in monocytes and macrophages in response to hypoxia and the high lactate conditions of the tumor microenvironment (Carmona-Fontaine et al., 2017; Colegio et al., 2014; Zhang et al., 2019).

Myeloid-derived suppressor cells (MDSC) are a functional heterogenous subset of myeloid cells that develop in response to cancer and other inflammatory conditions, and act as potent suppressors of T cell activity. MDSC are defined by a number of immunoregulatory molecules, including Arg1 (Cheng et al., 2008; Gallina et al., 2006; Huang et al., 2006; Kohanbash et al., 2013; Meyer et al., 2011; Sinha et al., 2008). However, our single-cell gene expression data did not detect significant numbers of *Arg1*-expressing cells that co-expressed *S100A9* and *Il4ra*, essential markers of MDSC (Bronte et al., 2016) (**Extended Data Fig. 5b**). Flow cytometric analysis confirmed our transcriptomic data, demonstrating no IL4R-expressing monocytic populations (**Extended Data Fig. 5c**). In the absence of these critical markers required for MDSC identification, these data suggest the tumorigenic effects of myeloid cells are due to TAM rather than MDSC.

Although Arg1 is consistently linked with malignancy, it is important to note that there have been no studies testing the role of Arg1 in a genetic model of cancer. To test the requirement of Arg1 to the development of NB, we used conditional deletion of *Arg1* in *Alk*^F1178L^; TH-MYCN mice. In this genetic strategy, the *Arg1*^flox^ allele was crossed to *Tek*-Cre mice (Tg(*Tek*-cre)^1Ywa^, referred to as Tie2-Cre hereafter) to generate a complete knockout in all hematopoietic cells and some endothelial cells. Our scGEX data and other cancer-related studies using implantable models demonstrate Arg1 is primarily expressed in myeloid cells (Arlauckas et al., 2018). As other known cells such as ILC2 and endothelia that express Arg1 were not detected by scGEX, nor by flow cytometric methods **(Extended Data Fig. 5d)**, we are confident that this genetic approach targets myeloid Arg1, as we have demonstrated previously (Kratochvill et al., 2015). The use of Tie2-Cre is also necessary to achieve complete deletion of the conditional Arg1 locus, as the commonly-used LysM-Cre (*Lyz2*-Cre) strain provides incomplete deletion (El Kasmi et al., 2008). Using this approach, we found that loss of myeloid Arg1 was protective against tumor formation (**Fig. 6e**). Pairwise comparisons demonstrate that Arg1 and CCR2 deficiency in *Alk*^F1178L^; TH-MYCN mice provide protection against tumor formation compared to wild-type mice, and whilst bone marrow-derived macrophages are required for tumorigenesis, conventional STAT6-mediated M2-polarized macrophages were not (**Fig. 6f**). Taken together, our findings indicate that a novel regulatory loop of Arg1-positive myeloid cells and non-antigen specific unconventional CD4^+^ promote a highly tumorigenic environment in this model of NB.

## Discussion

Our results indicate that a major pro-tumor effect of myeloid cells is through pathways that deplete amino acids. We rescued the lethal effects of *Alk*^F1178L^; TH-MYCN NB by depleting macrophages, CD4^+^ T cells, or myeloid Arg1; any one of these manipulations was sufficient to block tumor formation. By contrast, neither the STAT6-regulated pathways involved in M2 macrophage polarization, a pathway linked to tumor formation, nor CD8^+^ T cells were required in this model. We demonstrate the co-localization of CD4^+^ T cells and myeloid cells within the tumor microenvironment, suggesting functional relationships between these cell types. These findings were made possible by a highly penetrant model of NB, which combines tyrosine hydroxylase-driven MYCN amplification with a known activating mutation in the proto-oncogene *Alk*, resulting in tumors that replicate the gene expression profiles and immune infiltration patterns of human pediatric NB. Given the stark downregulation of MHC class I and ß2-microglobulin expression on both human and murine tumor cells, we anticipated NK cells would accelerate tumor formation, they are not required during these early stages of tumor formation and growth.

The adrenal medulla is populated with sparse numbers of macrophages derived from the fetal liver, but it is possible that some or all macrophages in the adrenal gland at the time of tumor formation may be bone marrow-derived. Without a clear genetic approach to eliminate embryonic-derived adrenal macrophages, we cannot determine their exact role in tumor formation. Through the use of *Ccr2*^-/-^ mice, which have substantial (but not complete) depletion of bone marrow-derived macrophage and monocyte populations, we determined that *Ccr2*^-/^; *Alk*^F1178L^; TH-MYCN mice had a trend to lower tumor incidence and significantly longer survival rates. Further work involving lineage-tracing would be necessary to determine the exact contribution of adrenal embryonic and bone marrow-derived macrophages to tumor formation. Nevertheless, we have demonstrated the importance of bone marrow macrophages to NB development and disease progression.

An important finding of the depletion studies is that CD4^+^ T cells, collectively, are pathogenic in NB, while B cells, CD8^+^ and NK cells are neutral for tumor formation and growth. This is supported by the finding that *Rag1*^-/-^; *Alk*^F1178L^; TH-MYCN mice, which lack an adaptive immune system, did not develop tumors. We speculate that an unconventional population of CD4 cells, potentially including Treg cells, are likely the pathogenic component of the CD4^+^ compartment. This requirement cannot be tested directly in our tumor model, since depletion of Treg cells either by antibodies or genetic ablation would result in the development of auto-immunopathology during the study window. Further studies are required to elucidate the exact mechanisms by which the deprivation of arginine triggers the development of these non-antigen specific unconventional cells, and precisely how these cells facilitate the tumorigenic process.

Our results suggest that inhibitors of Arg1 and other immune modulatory interventions may have significant value in cancer therapy. Immunotherapy thus far has been focused on promoting anti-tumor effects of NK or antigen-specific T cells. The paradigm suggested here is that disruption of the pro-tumoregenic, tissue forming effects of immune cells, including T cells and macrophages, may be therapeutically beneficial in NB, and perhaps in other low mutation burden developmental tumors.

## Online content

Extended data contains 6 figures and 5 tables.

## Acknowledgements

We thank the staff of the St. Jude Center for In Vivo Imaging and Therapeutics for their dedication and expertise, St. Jude Transgenic Core Unit for zygote and ES cell injections, Stefan Schattgen, Jessica Haverkamp, Dorian Obino, Crystal Neely and Lidija Barbaric for assistance, and the St. Jude Department of Immunology Flow Cytometry facility for cell sorting. This work was supported by the NIH grant CA189990, the Max Planck Gesellschaft, DFG Grants FOR 2599 and TRR 127, the Key For A Cure Foundation, the American Lebanese Syrian Associated Charities, and NIH Cancer Center P30 CA21765.

## Author contributions

L.V. performed the majority of the experiments, including mouse generation and characterization, breeding and genotyping, data acquisition and analysis, in vivo cellular depletions, flow cytometry, and in vitro assays. E.K.A. generated scGEX and spatial transcriptomics data, performed scGEX data analysis, and designed and generated figures. J.C.C. performed analyses on tumor incidence and survival, microarray expression, scGEX, spatial transcriptomics, TCGA expression data, and TCR repertoire, and generated figures. T.L.W. performed TCR repertoire experiments. C.G. performed the spatial transcriptomics sectioning and imaging. M.R. and L.Z. provided additional technical support. A.B. performed the human immunohistochemistry experiments. D.F. performed comparative transcriptome analysis between human and mouse NB. S.P. provided advice on CRISPR/Cas9 targeting strategies. S.S.C. provided advice and support for the animal studies. P.G.T. supervised the immunology experiments and contributed to data interpretation. P.J.M conceived the project and designed the *Alk* mutations. J.C.C. and E.K.A. edited the manuscript. L.V., P.G.T., and P.J.M. designed experiments, analyzed the data, and wrote the paper.

## Methods

### Mice

C57BL/6, *Ccr2*^-/-^ (B6.129S4-*Ccr2*^*tm1Ifc*^/J), *Stat6*^-/-^ (B6.129S2(C)-*Stat6*^*tm1Gru*^/J), *Rag1*^-/-^ (B6.129S7-*Rag1*^tmMom^/J) and OT-II (B6.Cg-Tg(TcraTcrb)425Cbn/J) mice were purchased from The Jackson Laboratory, and genotyped according to Jackson Laboratory protocols. *Arg1* floxed mice were crossed to Tie2-Cre. All founder animals were screened by PCR. For *Alk*^F1178L^ or *Alk*^Y1282S^ mutations, PCR products were digested with NsiI or EcoRV, respectively, to distinguish wild-type or mutant *Alk* alleles. All oligonucleotides are listed in **Extended Data Table 5**. All mice used in this study were bred and housed within a single cubicle within a larger room. Mice were maintained in a 12 hr day-night cycle, with constant temperature and humidity. Mice were bred using strategies to collect the greatest number of mice heterozygous for one *Alk* allele and TH-MYCN. Breeder mice bearing one mutant *Alk* allele have the same tumor penetrance as study mice, therefore loss of breeder mice was minimized by crossing males bearing one *Alk* mutant allele to multiple females bearing two wild-type *Alk* alleles. Mice were maintained by interbreeding to minimize space and the numbers of mice used. The genetic backgrounds of all mice used was mixed and originated from 129v/J (TH-MYCN) and C57BL/6. Following genotyping, mice were gender-segregated and assigned to imaging groups. Each animal on study was entered sequentially into a database. The St. Jude Children’s Research Hospital Institutional Animal Care and Use Committee approved all studies performed.

### Murine neuroblastoma study inclusion and exclusion criteria

Inclusion criteria were presence of the TH-MYCN allele, and heterozygosity for one *Alk* mutant allele and an age range from 1-3 weeks post-weaning. Either gender was used and noted. Criteria for analysis were the presence or absence of an ultrasound-detectable tumor (quantified by staff unrelated to this study) at any point within a 7 week period, where mice were imaged once per week. Endpoints were (i) an animal welfare issue warranting euthanasia at any point in the imaging period, (ii) a tumor of >500 mm^3^, (iii) a tumor whose growth rate indicates a size of >500m^3^ would be reached before the next imaging session, (iv) death, or (v) 7 consecutive imaging sessions. For final analysis and survival data, the age of all mice was averaged to 80 days to account for minor variations in days of age at the first imaging session. All raw data is available in **Extended Data Tables 1** and **2**.

### Ultrasound imaging

Fur was removed from the ventral side of each animal using Nair. Technicians in the St. Jude Center for In Vivo Imaging and Therapeutics performed ultrasound scanning on mice weekly using a VEVO-2100 and determined tumor volumes using VevoLAB 3.0.0 software. All ultrasound data were acquired in a blinded fashion.

### Immune cell isolation

Tumors were excised, mechanically dissociated and transferred to basal medium containing 0.1% collagenase Type IV and 150 mg/ml DNase I (Worthington), and incubated for 30 min shaking at 37°C. Tumors were then manually dissociated and passed through a 70 μm nylon mesh. Cells were centrifuged for 5 min at 400*g*, resuspended in PBS and overlaid on a 35%/60% Percoll gradient. Gradients were centrifuged at 2000*g* for 20 min at 4°C with no brake. Cells were collected from the 35%/60% interface and washed with PBS, prior to further analysis.

### Flow cytometry

Cells were treated with 1:100 dilution of CD16/CD32 blocking antibody (clone 2.4G2, Tonbo) and labeled with 1:500 Ghost Dye Violet 510 (Tonbo) according to manufacturer’s protocols. The following antibodies were used: B220 BV605 (BioLegend, clone RA3-6B2, lot# B248977, dilution 1:200), CD3 BV421 (BioLegend, clone 17A2, lot# B262915, dilution 1:200), CD3 BV605 (BioLegend, clone 17A2, lot# B233157, dilution 1:200), CD4 APC (BioLegend, clone RM4-5, lot# B261590, dilution 1:200), CD8a FITC (BioLegend, clone 53-6.7, lot# 8152865, dilution 1:200), CD8a BV785 (BioLegend, clone 53-6.7, lot# B266842, dilution 1:200), CD11b BV605 (BioLegend, clone M1/70, lot# B272198, dilution 1:200), CD11b BV785 (BioLegend, clone M1/70, lot# B253527, dilution 1:200), CD11b Pacific Blue (BioLegend, clone M1/70, lot# B211706, dilution 1:200), CD11c APC/Fire 750 (BioLegend, clone N418, lot# B250821, dilution 1:200), CD19 BV421 (BioLegend, clone 6D5, lot# B246776, dilution 1:200), CD25 PE (BioLegend, clone PC61, lot# B246222, dilution 1:200), CD45 APC/Fire 750 (BioLegend, clone 30-F11, lot# B260280, dilution 1:200), CD206 BV711 (BioLegend, clone C068C2, lot# B221988, dilution 1:200), CD273 PE (BioLegend, clone TY25, lot#B257232, dilution 1:200), CD274 APC (BioLegend, clone 10F.9G2, lot#B241508, dilution 1:200), CD276 AF647 (BioLegend, clone MIH32, BD, lot#8200854, dilution 1:200), F4/80 PE (BioLegend, clone BM8, lot# B251637, dilution 1:200), Gr1 BV605 (BioLegend, clone RB6-8C5, lot#B288302, dilution 1:200), GD2 PerCP/Cy5.5 (BD, clone 14G2a, lot#7200843, dilution 1:200), I-A/I-E PE (eBioscience, clone M5/114.15.2, lot# E01733-1633, dilution 1:200), IFN-gamma BV785 (BioLegend, clone XMG1.2, lot# B246604, dilution 1:100), Ly6C APC (BioLegend, clone HK1.4, lot# B204268, dilution 1:200), Ly6G FITC (BioLegend, clone 1A8, lot# B261238, dilution 1:200), Ly6G BV605 (BioLegend, clone 1A8, lot# B252817, dilution 1:200), Gr1 BV605 (BioLegend, clone RB6-805, lot# B288302, dilution 1:200), IL7RA PerCP/Cy5.5 (BioLegend, clone A7R34, lot# B249756, dilution 1:200), NK1.1 BV605 (BioLegend, clone PK136, lot# B261612, dilution 1:200), Sca-1 BV785 (BioLegend, clone D7, lot# B244413, dilution 1:200), ST2 PE/Dazzle 594 (BioLegend, clone DIH9, lot# B263231, dilution 1:200). To determine Foxp3 expression, cell surface stained cells were treated with Foxp3 Transcription Factor Staining Buffer Kit (Tonbo) according to manufacturer’s instructions, and stained with anti-Foxp3 AF488 (clone MF-14, BioLegend, lot# B248077, 1:50 dilution) or rat IgG2bϰ AF488 (clone RTK4530, BioLegend, lot# B211996, 1:50 dilution) isotype control. Data were acquired with a BD LSRFortessa X-20 and analyzed in FlowJo v10 (Treestar).

### Human immunohistochemistry

Patient sample analysis was approved by the institutional review board at St. Jude Children’s Research Hospital. Immunohistochemical studies were performed on archival formalin-fixed paraffin-embedded tissue blocks from 5 primary human NB (3 with *MYCN* amplification; 2 without *MYCN* amplification). A representative block of each tumor was sectioned at 4μm thickness and subjected to immunohistochemistry using the following panel of antibodies: CD68 (DAKO, clone PG-M1, lot# 20047721, 1:50 dilution), CD163 (Cell Marque, clone MRQ-26, lot# 13717, dilution 1:50), S100 (DAKO, rabbit polyclonal, lot# 20014502, dilution 1:8000); Arginase-1 (Cell Signaling, clone D4E3M, lot# 1, dilution 1:800).

### Antibodies

For in vivo CD4^+^ and CD8^+^ lymphocyte depletions, 0.5 mg purified and lipid absorbed CD4 ascites (Harlan Sprague-Dawley, clone GK1.5, lot# 200074), CD8 ascites (Harlan Sprague-Dawley, clone 2.43, lot# 70088) or vehicle control was administered intra-peritoneally to mice every 2 days until a study endpoint was reached. Depletion was confirmed by flow cytometric analysis using anti-CD4 clone RM4-5 or anti-CD8 clone 53-6.7. For in vivo NK cell depletions, mice were administered 100µg anti-NK1.1 antibody (BioXCell *InVivo*Plus, clone PK136, lot# 693118K3) or vehicle control every 4 days until endpoint was reached. Depletion was confirmed by flow cytometric analysis using anti-Nkp46 antibody. For in vivo B cell depletions, mice were administered 300µg of both anti-CD19 (BioXCell *InVivo*MAb, clone 1D3, lot# 701318J3) and anti-B220 (BioXCell *InVivo*Mab, clone RA3.3A1/6.1, lot# 754419A2) or vehicle control every 4 days until endpoint was reached. Depletion was confirmed by flow cytometric analysis using anti-CD19 clone 6D5. All depleting antibody lots were tested for endotoxin by the limulus amebocyte assay and found negative.

### TCR Transgenic CD4 Transfer

Spleens and lymph nodes were harvested from OT-II mice and manually dissociated through a 70µm nylon mesh. Cells were centrifuged for 5 min at 400*g*. After red cell lysis, cells were resuspended in PBS containing 0.5% FBS and 2mM EDTA pH8.0 and treated with CD16/CD32 blocking antibody. Cells were then surface stained with anti-CD4 and anti-Vβ5.1, 5.2 antibodies, and cell sorted. *Alk*^F1178L^; TH-MYCN were intravenously administered with 5 x 10^6^ CD4^+^ Vβ5.1, 5.2^+^ cells one week prior to study enrollment. After detection by ultrasound, tumors were processed as detailed above for downstream processing.

### Bulk transcriptome analysis

Human primary NB microarray data from GEO sources (GSE12460, GSE13136) were RMA normalized, summarized and batch corrected in Partek Genomics Suite 6.6 (St. Louis, MO). These data were merged to RMA summarized public human Ewing’s sarcoma (EWS), retinoblastoma (RB) and rhabdomysarcoma samples (RMS) (E-TABM-1202 ArrayExpress, GSE29683, GSE37372). The data was de-duplicated such that each gene symbol was represented by only one probeset. The probeset with the highest overall average expression across samples was retained. Means for each tumor class were calculated: NB MYCN amplified, non-amplified NB, EWS, RB, and RMS in STATA14/MP (College Station TX). A log fold change (FC) was calculated by subtracting the global mean of the non MYCN amplified tumor class means from the MYCN amplified NB: log FC = MYCN amplified NB – (non-amplified NB + RB + EWS + RMS)/4. Genes were the log FC exceeded 3 (equivalent to an eight-fold increase in MYCN B samples) were retained as signature genes. The human mean data for these select genes were then matched by gene symbols to de-duplicated mouse NB model data (GSE98763, GSE27516). The data for these 70 selected genes were then z transformed, hierarchically clustered, and a heatmap produced using Partek Genomics Suite 6.6.

### Single-cell gene expression

Tumor-bearing mice were injected intravenously with 10µg anti-CD45 APC to label circulating cells 15 minutes prior to harvest. Tumor was then processed and stained with a viability stain and anti-CD45 PE. After pre-gating on APC negative cells, live CD45^+^ PE and live CD45^-^ PE populations were sorted, centrifuged at 400*g* for 5 minutes, resuspended in 0.04% BSA in PBS. CD45^-^ and CD45^+^ populations were combined and counted by hemacytometer. Approximately 20,000 cells were loaded on the Chromium controller (10X Genomics) with the single-cell 3’ kit (V2) to capture approximately 10,000 cells. The remaining steps were performed according to the manufacturer’s protocol. Sequencing was performed on the Illumina Novaseq to generate 500M clusters per library. The raw reads were processed using Cell Ranger v3.1.0 with the mm10 (v3.0.0) reference. Feature matrixes were analyzed in Seurat (v3.2.0) (Stuart et al., 2019); briefly, cells with greater than the library-specific 98^th^ percentile of UMIs or genes, cells with fewer than 300 genes, and cells with greater than 10% mitochondrial expression were excluded before libraries were individually normalized via SCTransform, with the percent of mitochondrial expression regressed out. Libraries were also integrated using standard parameters before dimensionality reduction (tSNE with 50 principal components and an exaggeration factor of 30) and cluster identification. Transcriptional clusters were annotated manually by considering expression of various cell type-specific genes (**Extended Data Fig. 1c**) (Franzén et al., 2019), as well as with SingleR (Aran et al., 2019).

### Spatial transcriptomics

Two tumors between 100-200mm^3^ were harvested from distinct mice and snap-frozen in OCT using isopentane that was cooled in a liquid nitrogen bath. Tumors were cryosectioned at -22°C to 10 microns for use in Visium Spatial Transcriptomics kit (10X Genomics). We detected optimal tissue permeabilization at 12 minutes, and we sequenced libraries on the Illumina NovaSeq platform at 28×120bp. Capture images, which were extremely large, were reduced in resolution and manually rotated to better facilitate automatic fiducial marker identification. Data were processed using SpaceRanger (v1.0.0, 10X Genomics) with automatic alignment of fiducial markers and automatic tissue identification. Seurat (v3.2.0) was used for downstream analysis, where data were normalized using SCTransform and merged for comparison across sections. The merged data were used to detect specific genes enriched in areas where *Arg1* was expressed by comparing enzyme-positive areas to enzyme-negative areas using a Wilcoxon Rank Sum test. scGEX signatures were independently integrated with individual sections using anchors identified via PCA. As spatial transcriptomics data are not at single-cell resolution and can encompass multiple cells and cell types per spatial capture area, we considered the resulting probabilistic classification scores as approximate proportions of signal owing to each signature. Co-localization of distinct cell subsets was identified by assessing co-occurring classification scores from distinct subsets within the same capture area. To test for statistical differences in co-occurring signals, we categorized spatial areas as having a macrophage classification score of 0 (“no macs”) or a positive macrophage classification score (“with macs”) and conducted Wilcoxon Rank Sum tests on the classification scores of other signatures.

### TCR repertoire analyses

CD4^+^ CD25^+^ cells were sorted into 384 well plates, where a single cell is in an individual well, with index sorting enabled. Two columns of each 384 well plate was left unsorted to serve as a negative control. Plates were sealed and stored at -80°C until reverse transcription and subsequent PCR. Reverse transcription was completed using SuperScript VILO (Invitrogen, cat# 1175550, lot# 2167700) based on manufacturer’s protocol. TCR αβ cDNA were amplified and sequencing with previously described methods (Dash et al., 2011, 2015; Wang et al., 2012). TCR sequencing output was analyzed via TCRdist (Dash et al., 2017).

### Statistics

All mice included into the study were analyzed sequentially. No power analysis was performed before the study began because the rate of tumor formation in the mutant ALK backgrounds in the St. Jude vivarium was unknown at the initiation of the study. Therefore, measurement of tumor formation was longitudinal with corresponding analysis to estimate significance or non-significance in tumor formation over time. At the cessation of the study, all primary data was independently checked against the original database by two people and entered into GraphPad Prism v7 for survival curve comparison. The Log-rank (Mantel-Cox) test was used as the primary statistic to estimate significance or non-significance. Exact *P* values for curve comparisons are reported in the figures. Two-tailed *t* tests or Kruskal-Wallis tests corrected for multiple comparisons were used as appropriate. Additional survival analyses were conducted using Cox proportional hazard regression models in the R *survival* package after verification of model assumptions and visualized using *survminer*. The R *ggplot2* package was used for the customized forest plot after conducting pairwise comparisons; FDR was used to adjust for multiple comparisons within each study group, with error bars representing the standard error of the coefficients as determined by the respective CoxPH model.

## Data availability

Single-cell gene expression data and spatial transcriptomics data are accessioned in the Short Read Archive in association with BioProject PRJNA662418. Microarray files have been deposited in the Gene Expression Omnibus (accession GSE126024).

**Extended Data Table 1**. Tabulation of all mice used in the analyzed study groups

**Extended Data Table 2**. Mice excluded from the study

**Extended Data Table 3**. Human NB ‘signature’ genes

**Extended Data Table 4**. Microarray data referring to Fig. 1.

**Extended Data Table 5**. Oligonucleotides used in this study.

**Extended Data Fig. 1**. Construction and characterization of a genetic model of NB. **a**, Mutagenesis of *Alk* by CRISPR/Cas9. Representative sequence analysis of the wild-type and *Alk*^Y1282S^ allele showing the Tyr>Ser mutation. * indicates the addition of silent point mutations introduced to reduce subsequent Cas9-mediated DNA cleavage. **b**, Representative sequence analysis of the *Alk*^F1178L^ allele. The introduced NsiI site to facilitate genotyping is indicated. **c**, t-SNE dimensionality reduction of mouse *Alk*^F1178L^; TH-MYCN tumor CD45^-^ and CD45^+^ cells based on single-cell gene expression (scGEX) data, with transcriptional clusters numbered for identification from most to least prevalent. **d**, Feature plots showing expression of a subset of genes known to be involved in NB tumors and cancer biology in general. **e**, Feature plots showing expression of a subset of genes utilized for annotation of major cell types.

**Extended Data Fig. 2**. NB downregulates MHC class I. **a**, Feature plot showing *B2m* expression in an *Alk*^F1178L^; TH-MYCN tumor, as identified by scGEX, demonstrating negligible expression in CD45^-^ cells. **b**, NB have relatively low expression of *B2M* compared to other pediatric malignancies. **c**, Incidence and survival curves in *Alk*^F1178L^; TH-MYCN mice treated with anti-NK1.1 antibody or control. Data were analyzed by the Log-rank test.

**Extended Data Fig. 3**. CD8^+^ Tregs are not present under normal conditions. Unlike tumors that developed in *Alk*^F1178L^; TH-MYCN mice that were depleted of CD4^+^ cells, control tumors do not contain significant numbers of CD8^+^ CD25^+^ Foxp3^+^ cells. Data are representative of *n*=2 tumors.

**Extended Data Fig. 4**. Lymphocyte-macrophage co-occurrence dynamics vary by cell subtype. **a**, Loess regressions of immune cell type classification scores as a function of macrophage classification scores. Plotted data include all spatial transcriptomics capture spots (n = 6,729) obtained from four independent tumor sections, each of which was analyzed independently. Shaded area is representative of the 95% confidence interval. **b**, Tumors from mice deficient in CCR2 are depleted of monocyte and macrophages populations. *Alk*^Y1282S^; TH-MYCN; *Ccr2*^-/-^ mice have fewer CD11b^+^ Ly6G^-^ Ly6C^+^ circulating monocytes compared to their wild-type counterparts. **c**, The flow cytometry gating strategy for myeloid populations is shown, as well as cytospins from sorted fractions. Representative flow plots from tumors originating in *Alk*^F1178L^; TH-MYCN; *Ccr2*^-/-^ demonstrate fewer infiltrating monocytes and macrophages compared to their wild-type counterparts. Data are representative of two independent samples. **d**, Percentages of mature macrophages (C fraction) in CCR2-deficient tumors from the CD11b^+^ population. Mice with *Alk*^F1178L^; TH-MYCN or *Alk*^Y1282S^; TH-MYCN background were combined. In **a** and **c**, the bars represent the mean of the samples with the SEM. *P* values were calculated by two-tailed *t*-test.

**Extended Data Fig. 5**. Myeloid cells, and not tumor cells, express Arg1. **a**, Human ARG1 cells in NB are consistent with sub-populations of myeloid cells that intermingle with CD68^+^, CD163^+^ cells but not dendritic cells (S100^+^). **b**, Profiling of Arg1^+^ cells reveals a TAM-like phenotype, rather than a MDSC phenotype. Feature plots from *Alk*^F1178L^; TH-MYCN myeloid cells showing expression of *Arg1* (far left), *S100a9* (middle left, upper) and *Il4ra* (middle left, lower). *Arg1*^*+*^ myeloid cells show minimal co-expression (yellow) with either *S100a9* or *Il4ra*, which are critical to their identification of MDSC. **c**, Flow cytometric analysis from CD45^+^ CD11b^+^ Ly6G^-^ Ly6C^+^ cells isolated from *Alk*^F1178L^; TH-MYCN tumors demonstrate minimal expression of IL4RA (CD124). **d**, ILC2 are not found within the tumor microenvironment. Lineage negative (CD3, B220, CD11b, Gr1, NK1.1) cells were assessed for the presence of ILC2, which express the surface markers Sca-1, IL7RA and ST2.

