## Supplementary figures and images for "Neuroblastoma formation requires unconventional CD4 T cells and myeloid amino acid metabolism"

### Extended Figures

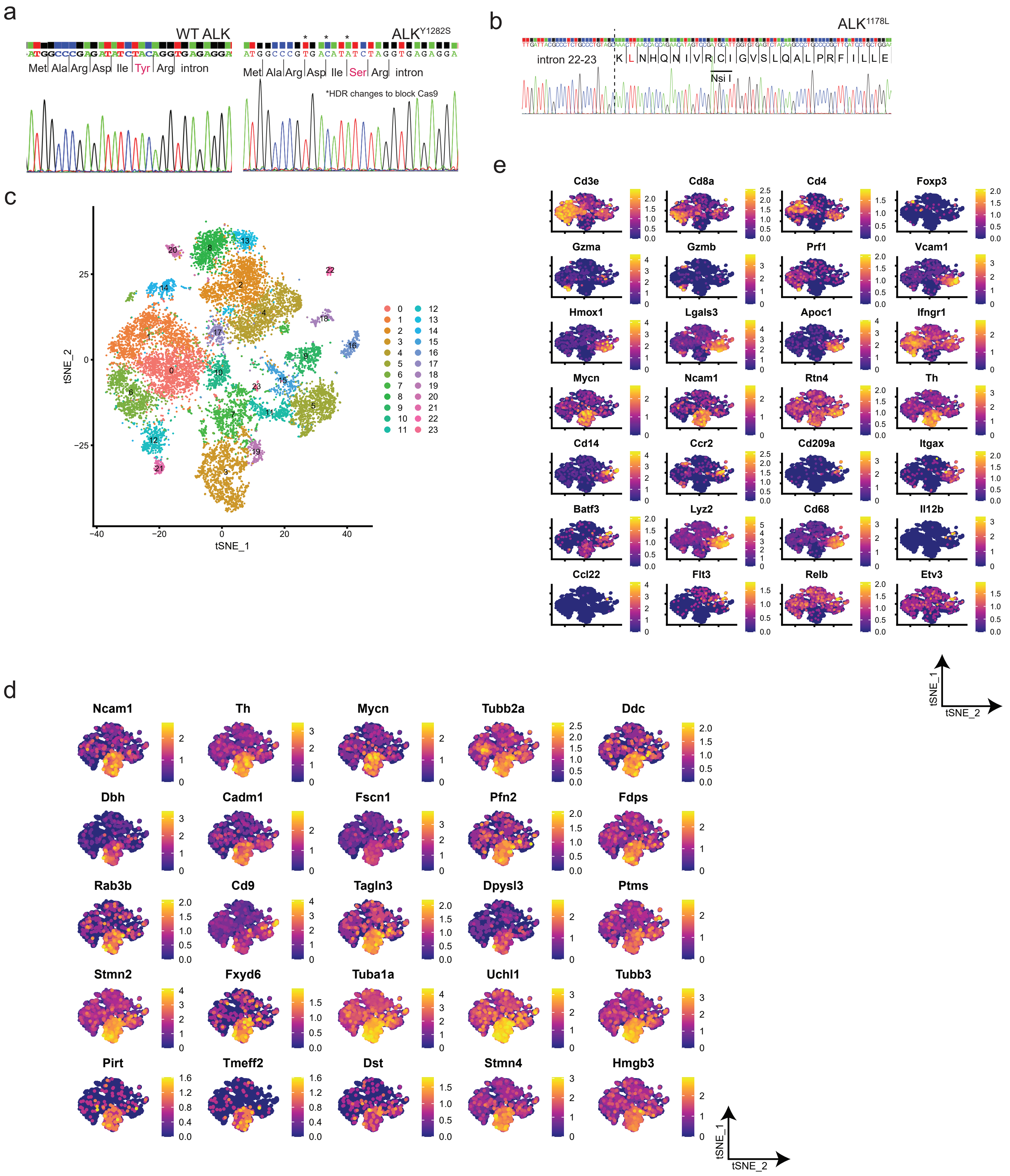


Extended Figure 1

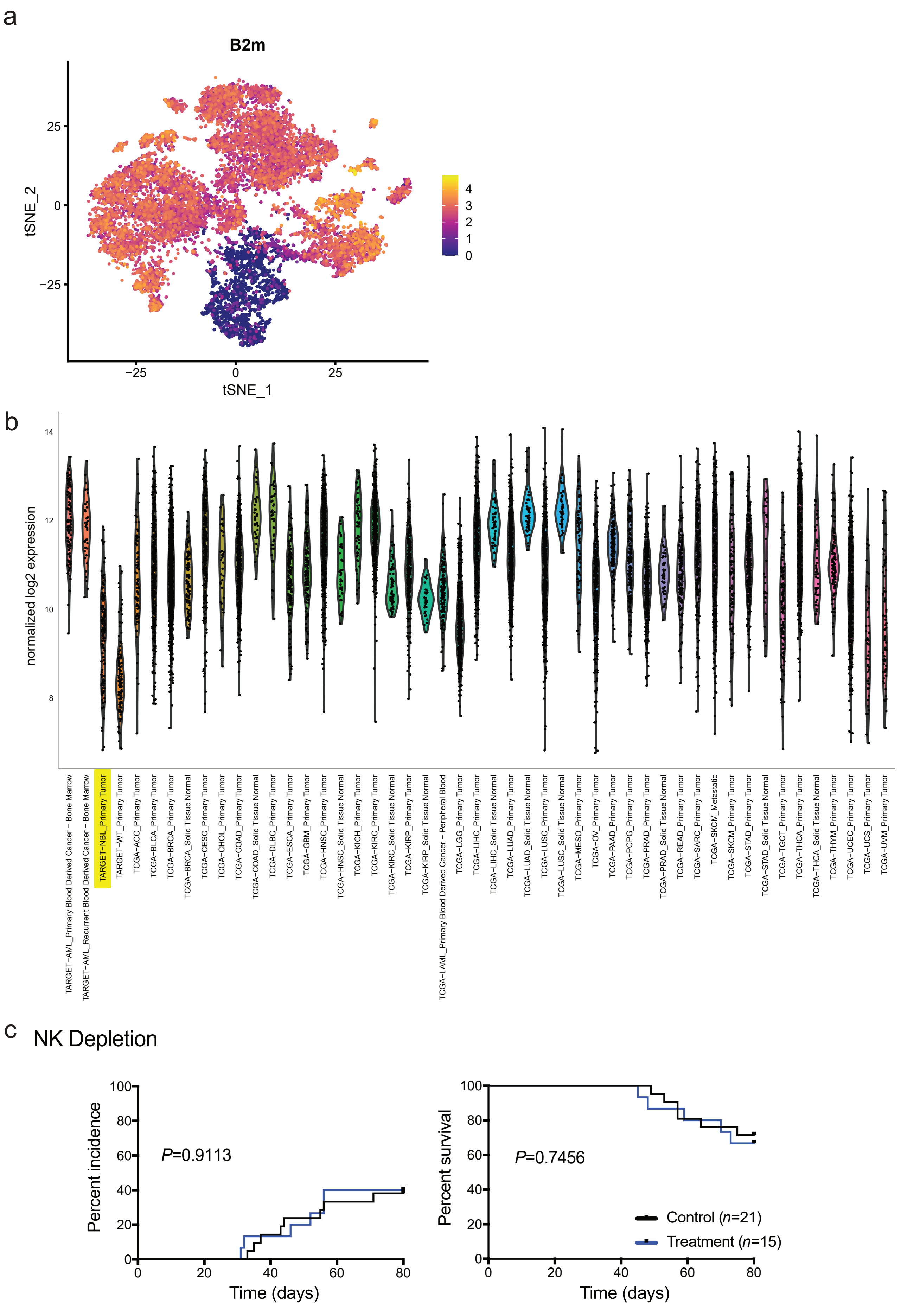


Extended Figure 2

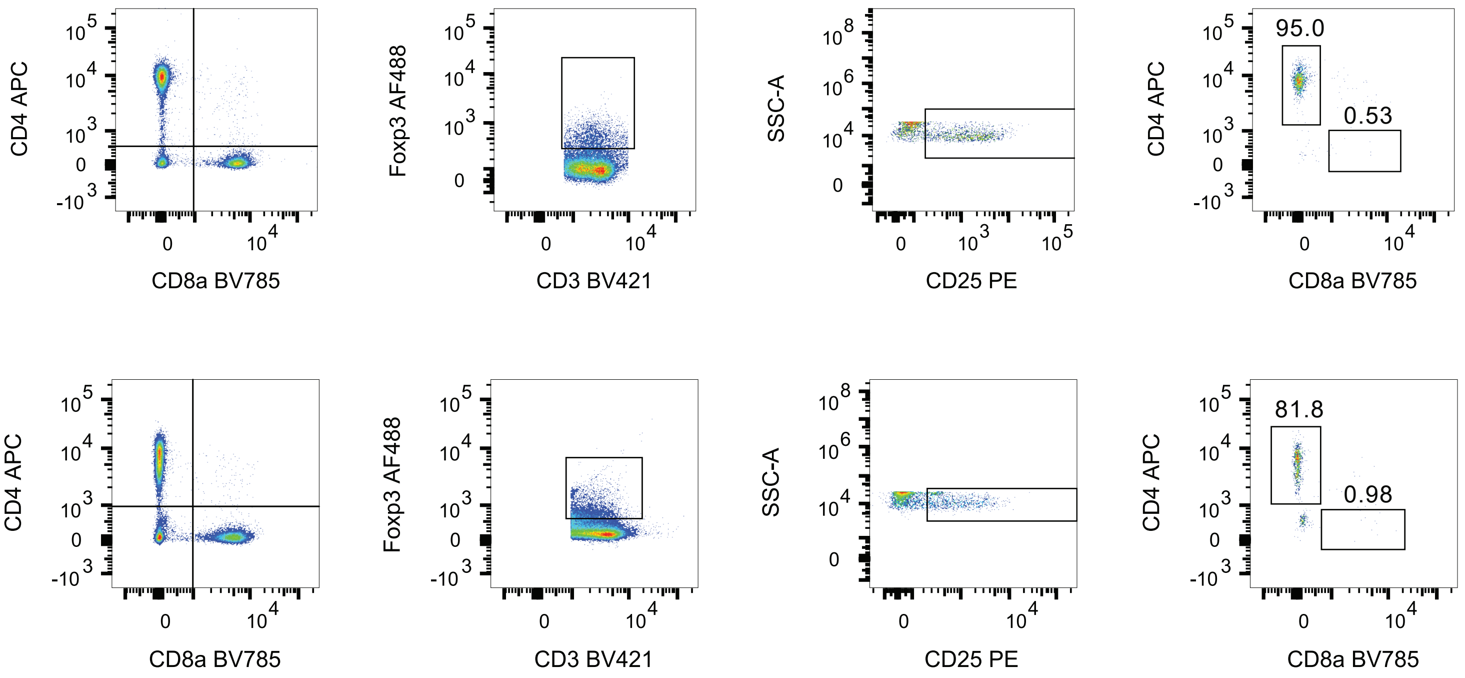


Extended Figure 3

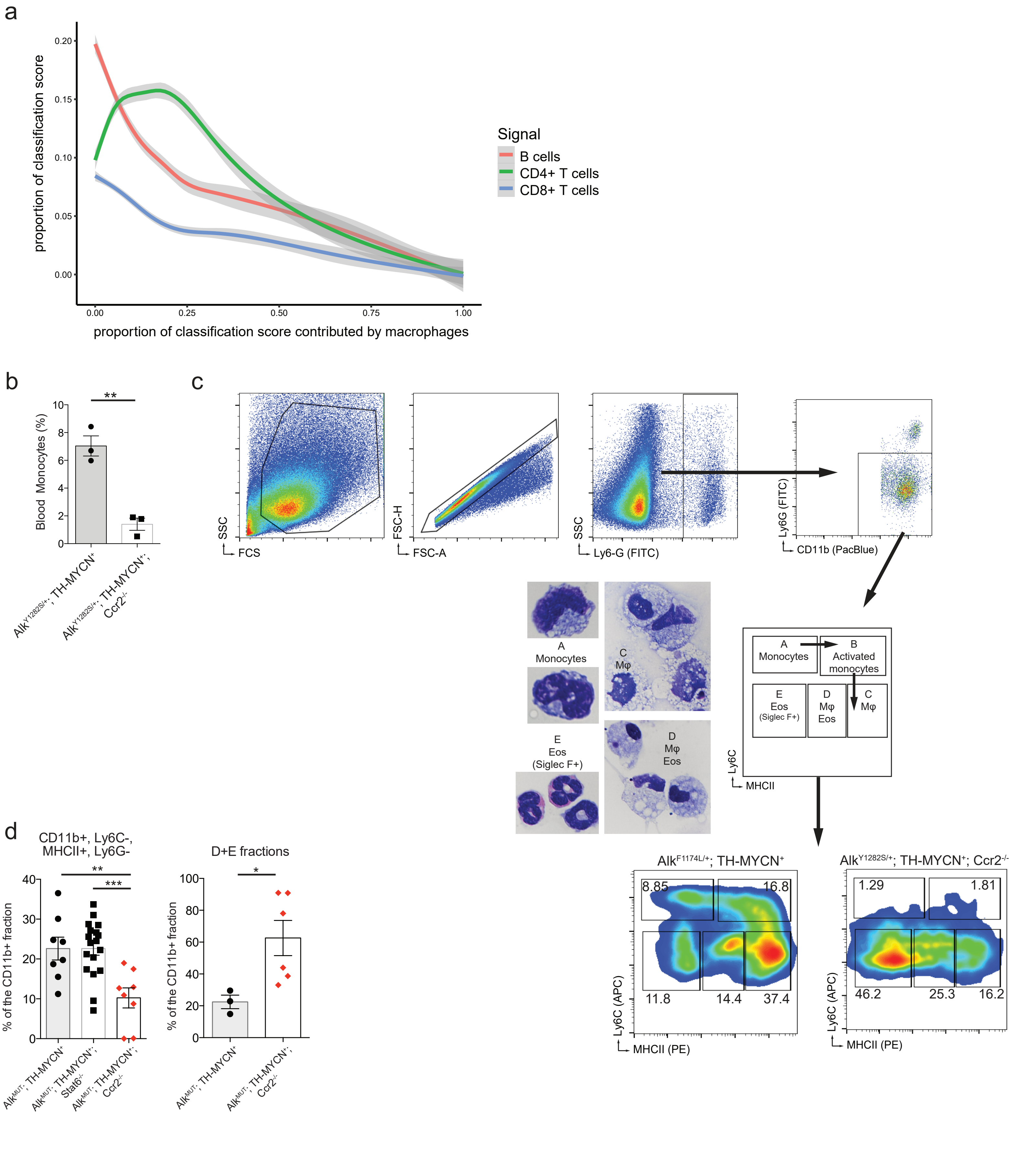


Extended Figure 4

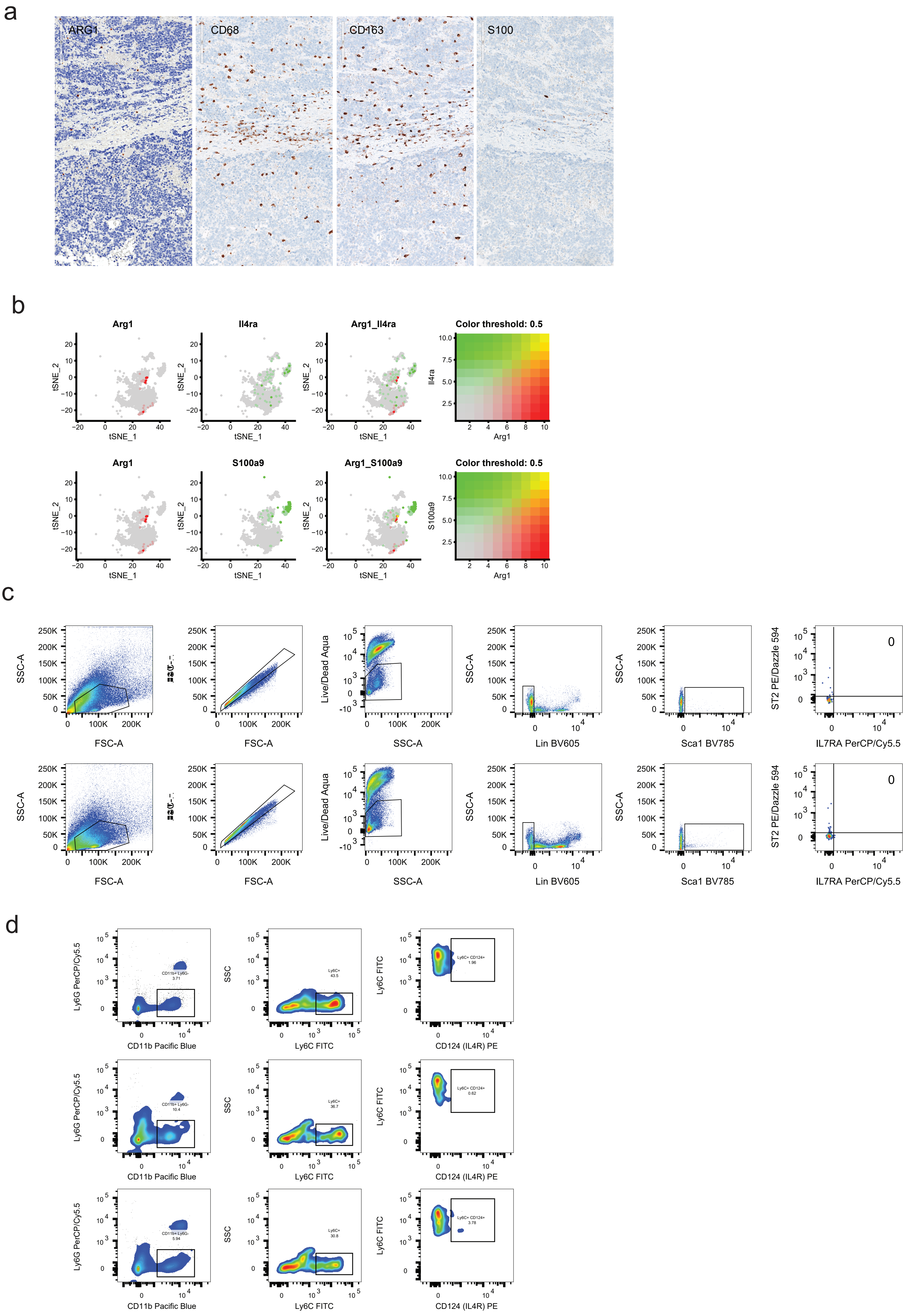


Extended Figure 5
