## Extended Data Table 5 for "Neuroblastoma formation requires unconventional CD4 T cells and myeloid amino acid metabolism"

Alk Y1282S
gRNA
TAATACGACTCACTATAGGGGATGGCCCGAGATATCTACGTTTTAGAGCTAGAAATAGCA
HDR Oligo CCAGGGATATTGCTGCTAGAAACTGTCTGTTGACCTGCCCAGGAGCTGGAAGAATAGCAAAGATTGGA GACTTTGGGATGGCCCGTGACATATCTAGGTGAGAGGATTGCCTTGCTCTGCCCACACCTTCCTCCACCA GCCAGTCTCCTCAAATAGATGCCTGGGCTGGCTTATATCTAATATCACCTTGTATCTTGTC
Genotyping Primers
Forward ACAGGGTGAAGGAAAGAACTGT
Reverse AGCCAAGGCAAACCATGCTA

Alk F1178L
gRNA
TAATACGACTCACTATAGGGAATATTGTTCGCTGCATCGGTTTTAGAGCTAGAAATAGCA
HDR Oligo CACAGCGTGATTGCTGAAGCCATGACTCCGTGCCCTGCTTCTCTTGATGGTCCCAGGACGGGCTCAGTTA AATTTGATTACGCCCTCTGCCCTGTAGCAAACTTAACCACCAGAACATAGTCCGATGCATTGGTGTGAGT CTACAAGCCCTGCCCCGCTTCATCCTGCTGGAACTCATGGCTGGCGGAGACCTCAAGTCC
Genotyping Primers
AGACCCCTCCACCCATATCC
ATGACCAGACCACAGGCTGA
